# *Aedes aegypti* aminopeptidase N3 is a functional binding receptor of *Bacillus thuringiensis* subsp. *israelensis* Cry4Ba toxin

**DOI:** 10.1101/2025.03.07.641997

**Authors:** Xiaozhen Yang, Wanting Huang, Jiajia Wei, Xiaoxuan Xu, Jackson Champer, Junxiang Wang

## Abstract

*Bacillus thuringiensis* is widely employed for biological control. It can effectively suppress populations of various mosquito species, including *Aedes aegypti*. However, the precise mechanism underlying the action of cry toxin secreted by *Bacillus thuringiensis* on *Ae. aegypti* remains elusive. In this study, we investigated one of the binding receptors of cry toxin, aminopeptidase N. Through comprehensive bioinformatics analysis involving whole-genome screening, genetic mapping, structural characterization, phylogenetic analysis, and spatiotemporal expression profiling, we identified twenty-nine homologs of *Ae. aegypti* aminopeptidase N. Further, we successfully expressed GST-APN3 protein in *E. coli* and demonstrated through ligand blot and ELISA assays that APN3 exhibits high affinity binding to Cry4Ba toxin (Kd = 20.53 nM). To elucidate the functional role of APN3 as a receptor mediating Cry4Ba activity in *Ae. aegypti* midgut cells (some of which express this gene at high levels), CRISPR/Cas9 technology was employed to knock out APN3. Our bioassay results revealed that APN3 knockout mosquito larvae had 2.9 to 4.1-fold higher resistance against Cry4Ba, indicating its crucial involvement as an active receptor mediating Cry4Ba activity. Overall, this study provides a foundation for elucidating the specific larvicidal mechanisms of Bt against mosquito populations.

## 1 Introduction

*Aedes aegypti* is the principal arthropod vector responsible for the transmission of several medically significant arboviral infections, most notably dengue fever^1,2^. Recent epidemiological data from the World Health Organization (WHO) reveal that approximately 4.1 billion people across 132 endemic countries are at substantial risk of dengue virus exposure^3^. The absence of universally approved prophylactic vaccines underscores the critical necessity for implementing effective vector control to disrupt disease transmission. Under the guidance of WHO, strategies such as “Integrated and Sustainable Vector Management” have yielded improvements in mosquito control^3,4^. Presently, biological control has gained international recognition as a preferred measure for disease and pest prevention and control. *Bacillus thuringiensis* (Bt), has emerged as the most extensively researched and widely applied biocontrol microorganism with its high insecticidal activity, broad spectrum efficacy, strong target specificity, and non-toxicity towards humans and animals^5–7^.

Bt synthesizes a diverse of proteins exhibiting insecticidal properties throughout its vegetative, growth, and sporulation phases. The primary insecticidal protein is known as crystal protein (Cry) toxin, while certain strains also produce cytolytic proteins (Cyt toxin) and vegetative insecticidal protein (VIP)^7,8^. Upon ingestion by the target insect, the Cry prototoxin undergoes dissolution in the alkaline environment of the intestine and subsequent activation through protease hydrolysis^8^. This process involves removal of nontoxic regions from both N- and C-termini to generate 3d-Cry active monomers. These monomers traverse across the peritrophic membrane surrounding the intestinal lumen and interact with receptor proteins on brush border membrane vesicles of epithelial cells^8^. Binding interactions between these receptors result in extensive epithelial cell death, leading to symptoms such as intestinal ulceration or sepsis ultimately causing mortality in insects. Based on findings from various *in vitro* and *in vivo* experiments conducted on different insects, it is generally believed that aminopeptidase N (APN), alkaline phosphatase (ALP), cadherin-like (CAD), and ATP-binding cassette transporter subfamily C are among the main receptor proteins involved^7^.

APN is a widely distributed ectopeptidase protein belonging to the M1 zinc metalloprotease family, found in animals, plants, and microorganisms. In insects, GPI-anchored APN membrane proteins serve as crucial binding receptors for Cry toxins^9^. *In vitro* binding experiments have demonstrated that APN proteins from various insects can bind to different Cry toxins, including *Manduca sexta*^10^, *Trichoplusia ni*^11^, *Anopheles quadrimaculatus*^12^, *Anopheles gambiae*^13^, *Ae. aegypti*^14–16^, and other Lepidoptera^17–20^ and Coleoptera^21^ insects. However, recent studies utilizing CRISPR/Cas9-mediated knockout of certain APN genes have revealed no alteration in sensitivity towards corresponding Cry toxins^22–24^. Therefore, further validation is required to elucidate the significance of APN in mediating the mechanism of action of Bt.

*Bacillus thuringiensis* subsp. *israelensis* (Bti) secretes a range of proteins that specifically target mosquito larvae, including Cry4Aa, Cry4Ba, Cry10Aa, Cry11Aa, Cyt1Aa, Cyt1Ca, and Cyt2Ba^25^. Among these proteins, Cry4Ba and Cry11Aa have been found to exhibit activity against *Ae. aegypti* larvae^25^. However, the precise role of APN in the mechanism of action of these two proteins on *Ae. aegypti* larvae remains unclear. In our previous study utilizing GST-Pull down and Co-IP techniques from *Ae. aegypti* brush border membrane vesicles, we identified five APN proteins that interact with either Cry4Ba or Cry11Aa^14^. Knocking out APN1 and APN2 did not impact the insecticidal activity of Cry4Ba and Cry11Aa^22^. To investigate whether other APN genes are involved in this process, we initially characterized the genomic structure features of APNs in *Ae. aegypti* through whole-genome identification as well as transcriptomic analysis of APN transcripts at different developmental stages and in various tissues. We then focused on investigating an interaction between an APN3 protein and Cry11Aa using brush border membrane vesicles from *Ae. aegypti*. The binding affinity between GST-APN3 fusion protein expressed in prokaryotic cells and Cry4Ba was determined through ligand blotting, and ELISA was performed to determine their binding affinity. Finally, CRISPR/Cas9 was employed to construct an APN3-KO gene knockout strain, confirming that APN3 functions as the functional receptor mediating low-dose activity of Cry4Ba.

## 2 Materials and Methods

### 2.1 Mosquito Strains and Rearing

The wild type strain of *Ae. aegypti* (Haikou strain) was provided by the Fujian International Travel Health Care Center (Fujian, Fuzhou, China). The EXU strain is a Cas9 knock-in strain derived from the wild-type strain through microinjection^26^. Both strains were reared for over 70 generations and 10 generations respectively without exposure to Bt toxins. All *Ae. aegypti* strains were maintained under controlled conditions at 26 ± 1 ℃, with a relative humidity of 83% ± 3% and a photoperiod of 14:10 h (light: dark). *Ae. aegypti* larvae were fed goldfish feed (Tetra, Germany), while adult mosquitoes received a diet consisting of a 10% sucrose solution. Moreover, pregnant female mosquitoes were provided with sterile defibrinated bovine blood as their nutritional source for oviposition.

### 2.2 Bt Strain and Purification of Cry4Ba and Cry11Aa Protoxins

The Bt strains individually producing recombinant Cry4Ba protein (pCG6- Cry4Ba) and Cry11Aa protein (pCG6-Cry11Aa) were provided by the Dr. Sarjeet R Gill’s Laboratory, University of California Riverside. The Bt bacteria were cultured in 1/2 LB (Luria-Bertani) medium supplemented with erythromycin at a final concentration of 12.5 μg/mL, and incubated at 30 °C. Once the crystal inclusions were fully released, the culture medium was removed and the pellet were washed three times by 1 M NaCl plus 0.03% Triton X-100, followed by distilled water rinses. The Cry4Ba was purified by isoelectric point precipitation method as described previously^27^, and protoxin was solubilized in alkaline buffer (50 mM Na2CO3/NaHCO3, pH 10.5). For Cry11Aa purification, density gradient centrifugation using sucrose solution was performed according to previous protocols^28^, followed by solubilization of protoxin in distilled water.

### 2.3 Sequence analysis of *AeAPN*

The genomic data of *Ae. aegypti* (AaegL5.2), *Culex quinquefasciatus* (CpipJ2.4) and *An. gambiae* (AgamP4.12) were downloaded from VectorBase database (https://www.vectorbase.org). The R language package biomaRt was utilized to extract the DNA sequences, cDNA sequences and amino acid sequences of APN gene family from three genomes. Additionally, relevant gene annotation information was extracted from both VectorBase and NCBI databases (https://www.ncbi.nlm.nih.gov). To further investigate the function of APN, the genomic structure of APN genes were visualized using MapGene2Chrom V2.1 (http://mg2c.iask.in/mg2c%5Fv2.1/), while features of amino acid sequence were analyzed by SMART (http://smart.embl-heidelberg.de). SignalIP 5.0 Server (http://www.cbs.dtu.dk/services/SignalP/) was applied for predicting the N-terminal signal peptide of APN, whereas transmembrane domain analysis of APN was conducted using TMHMM Server v. 2.0 (http://www.cbs.dtu.dk/services/TMHMM/). Furthermore, GPI-anchored points of APN were predicted utilizing big-PI Predictor (https://mendel.imp.ac.at/gpi/gpi_server.html), PredGPI predictor (http://gpcr2.biocomp.unibo.it/gpipe/pred.htm) and KohGPI (http://gpi.unibe.ch). Finally, NetNGlyc 1.0 Server (http://www.cbs.dtu.dk/services/NetNGlyc/) and NetOGlyc 4.0 Server (http://www.cbs.dtu.dk/services/NetOGlyc/). were employed for predicting N-glycosylation sites and O-glycosylation sites in APN.

The ClustalW program of MAGAX 10.1.7 was applied to perform multiple sequence alignment for the APN gene family of *Ae. aegypti*, while Neighbor-Joining was utilized for evolutionary tree clustering analysis selecting the default parameter. The conserved domains, motifs, and protein sequence conduction were visualised using TBtools software. Additionally, further phylogenetic analysis of APN gene families in three pathogenic mosquitoes was performed, and the resulting evolutionary tree was modified using iTOL (https://itol.embl.de).

### 2.4 Spatio-Temporal Expression Patterns of APNs in *Ae. aegypti*

The transcriptome data of *Ae. aegypti* at different developmental ages were obtained from Matthews et al. ’s study, including 1st to 4th instar larva, pupa, and male and female adults. RNA was extracted separately from the alimentary tracts (ATs) and bodies minus alimentary tracts (BMATs) of twenty 4th instar larvae, followed by transcriptome sequencing conducted by Biomarker Technologies company (data not yet published). The temporal and spatial expression patterns of the APN gene in *Ae. aegypti* was classified and mapped using TBtools software.

### 2.5 Prokaryotic expression of *AeAPN3* protein

The synthesized cDNA was used as a template to amplify AeAPN3 CDS sequences with specific primers (Table S1). The resulting PCR products were then cloned into the pCold-GST vector using the In-Fusion Snap Assembly Master Mix (Takara, Japan) according to the manufacturer’s instructions. Subsequently, assembly product was transformed into E. coli JM109 cells (Takara, Japan). For protein expression, the pColdAeAPN3 recombinant plasmid was further transfected into E. coli BL21 (DE3) cells (Takara, Japan). Purification of AeAPN3 protein was achieved through Glutathione Sepharose 4B chromatographic column. To confirm the expression of recombinant AeAPN3 protein, Western blot analysis was performed using an anti-GST antibody (TransGen, China).

### 2.6 Binding Analysis of AeAPN3 Protein to Cry4Ba Toxin

Ligand blot assays and Binding enzyme-linked immunosorbent assays (ELISAs) were conducted to assess the binding affinity between AeAPN3 protein and Cry4Ba toxin. For ligand blot assays, equal amount of GST-APN3 protein, GST tag protein and Cry4Ba crystal protein were separately separated by SDS-PAGE gel electrophoresis and transferred onto a PVDF membrane by wet transfer system. After overnight blocking at 4 ℃ in PBST buffer containing 5% BSA, the Cry4Ba protoxin (Cry4Ba: TBST=1:150) was added to the above blocking buffer at 25 ℃ for 2 h. After incubation, the membrane was washed three times with PBST for15 min each time. Then the membrane was hybridized with the primary antibody anti-Cry4Ba antiserum (adding to the blocking buffer at 1:3000 dilution) at 25 ℃ for 50 rpm 2 h, followed by three washes and subsequent incubation with alkaline phosphatase (AP) -conjugated secondary antibody (1:1000) for 2 h. After another round of washing three times, the membrane was visualized with BCIP/NBT Alkaline Phosphatase Color Development Kit.

For ELISAs, each well of the 96-well ELISA plates was coated overnight with 5 μg AeAPN3 protein in a final volume of 100 μl of PBS at 4 ℃ and blocked by blocking buffer at37 ℃ for 2 h. The plate was washed 3 times with 200 μl of PBST. Then, different concentrations, ranging from 0 to 320 nM, of Cry4Ba protoxins were independently added to wells in a final volume of 100 μl of blocking buffer. After incubated at 37 ℃ for 1 h, the plate was washed three times with PBST, and treated with primary antibody anti-Cry4Ba antiserum (adding to the blocking buffer at a dilution ratio of 1:10000) for 1 h at 37 ℃. Following this step, the plate was washed three times using TBST, and incubated with a horseradish peroxidase (HRP) -conjugated streptavidin antisera (1:3000) for 1 h. After three times washing by TBST, a volume of 100 μL of TMB was added to each well and incubated in darkness for 10 min at 37 °C. Finally, 50 μL/well of 2 M H2SO4 was added to terminate the reaction, and the plate was sent for colorimetry to measure the absorbance at 450 nm using microplate reader. The ELISA-binding plots were generated in GraphPad prism 8.0 and the Kd values were calculated using Curve Expert 1.4.

### 2.7 Embryo Microinjection and Screening of Homozygous Strain

The CRISPR/Cas9 target sites were designed in the fourth exon of AeAPN3 genes using CRISPOR program (http://crispor.tefor.net/). Potential off-target effects were evaluated by Cas-OFFinder (http://www.rgenome.net/cas-offinder/). The sgRNA was synthesized and purified in vitro using specific primers (Table S1). The Cas9 protein and sgRNA were microinjected into fresh exuCas9-kmo embryos^26^. Genomic DNA was extracted from each individual mosquito by leg pulling to identify gene mutants until a homozygous strain (APN3- KO with DsRed+ KMO-) was obtained.

Then, phenotypic observation, fluorescent screening and gene sequencing of APN3-KO with DsRed+ KMO- strain were performed to select individuals with non-fluorescent and normal eyes for establishing a new APN3 gene knockout homozygous line (APN3-KO), excluding any potential influence of endogenous expression of the CRISPR/Cas9 system and deletion of kmo gene on subsequent functional verification. More details as described before^22^. In the subsequent breeding of the line, 5 Aedes mosquitoes were randomly selected for mixed PCR and sequencing to ensure the genetic homozygosity of the knock out line.

### 2.8 Bioassay of Bti Cry Toxins

The susceptibility to Bti Cry4Ba and Cry11Aa toxins was determined for different strains of Ae. aegypti as previously described. The wild-type and APN3-KO strain of *Ae. aegypti* were fed to the beginning of age Ⅳ, and 25 larvae were transferred to 20 ml filtered water. Then Cry4Ba or Cry11Aa crystal proteins were added at concentrations according to previous experiments. At least 5 concentration gradients were set, with three biological replicates performed. Larval mortality was calculated 24 h later, and the lethal concentration required to kill 50% of larvae (LC50) was calculated using PoloPlus software.

## 3 Results

### 3.1 Chromosome localization of APN members in *Ae. aegypti*

A total of 29 APNs were identified from the *Ae. aegypti* genome based on the conserved APN core motif. Chromosomal localization analysis revealed that 26 *AeAPN* genes were distributed on three chromosomes (1, 2, and 3). However, *APN4* (AAEL005821), AAEL020609, and AAEL022733 were mapped to unassigned genomic scaffolds (NIGP01002224, NIGP01000461, and NIGP01001046), respectively. Spatial distribution analysis indicates chromosomal preference, with 73.07% (19/26) of chromosomally anchored APN genes residing on chromosome 1. These genes are predominantly clustered in the subtelomeric regions or nearby. The APN genes of *Ae. aegypti* predominantly occur in the form of gene clusters. For instance, *APN1* (AAEL012778), *APN3* (AAEL012774), and AAEL012776 are arranged consecutively on chromosomes 1. Similarly, on chromosome 3, there is a gene cluster consisting of *APN2* (AAEL019828), AAEL019829, AAEL008162, and AAEL008163. This clustered genomic organization indicates that the APN gene family of *Ae. aegypti* has undergone multiple evolutionary replication events (Fig. 1).

**Fig. 1.**
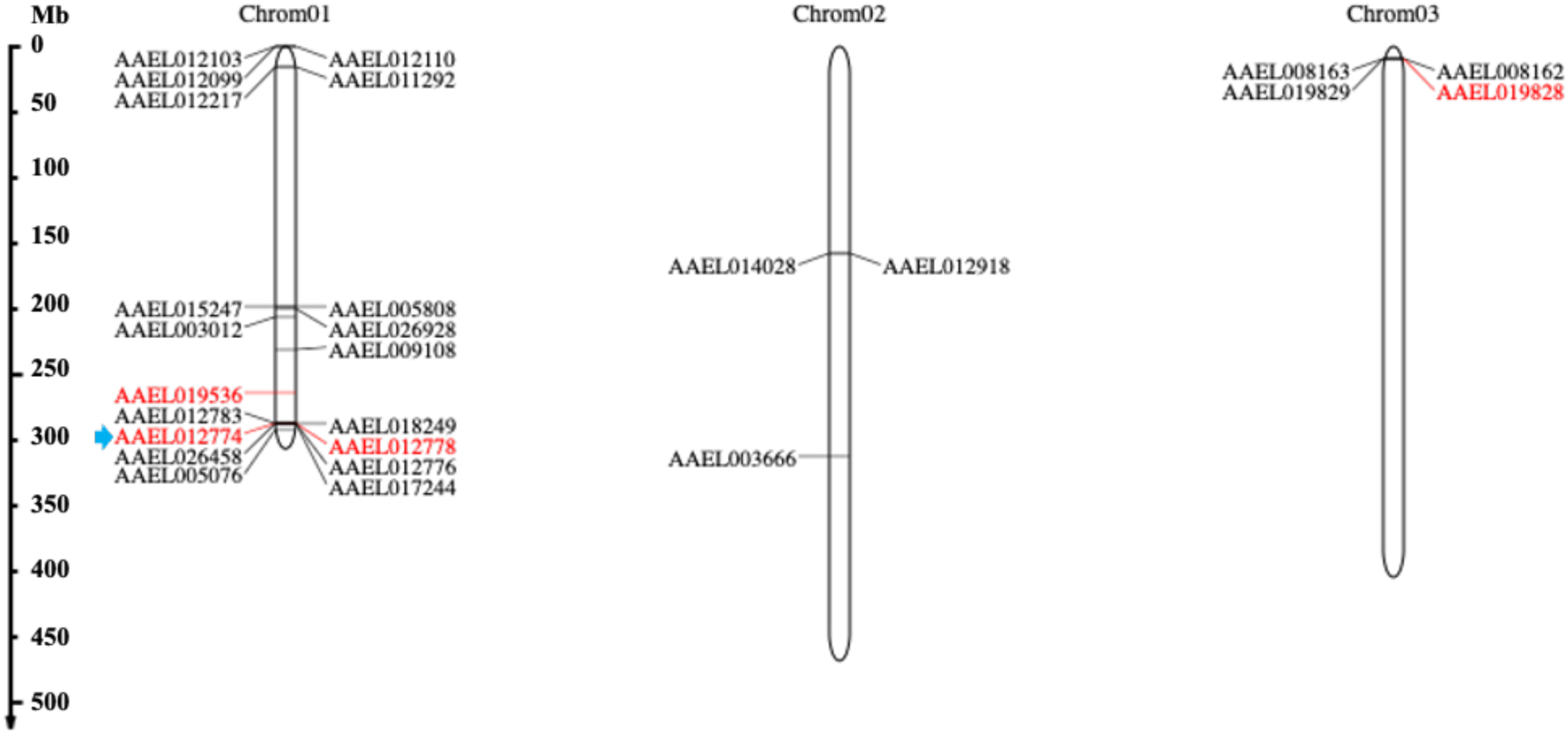
**Physical map of APN genes in *Ae. Aegypti* chromosomes**. Cry-binding APN isoforms are marked in red. The blue arrow indicates the location of *AeAPN3*.

### 3.2 Gene structures and conserved motifs analysis of AeAPNs

Bioinformatic characterization of AeAPN protein structures revealed that 11 AeAPNs have predicted GPI anchoring sites at the C-terminal, including four Bti-binding proteins (APN1, APN2, APN3, and APN5 (AAEL019536)). Comparative analysis of N-terminal modifications demonstrated commonalities and variations: while all five AeAPNs (APN1-5) possessed identifiable signal peptides and N-glycosylation sites, only four had O-glycosylation sites. APN1 exhibited a unique absence of O-glycosylation modification. Notably, APN5 emerged as the most extensively glycosylated isoform, containing 7 N-linked and 13 O-linked glycosylation sites (Table S2). Phylogenetic reconstruction of *Ae. aegypti* APNs revealed substantial sequence divergence across family members, indicative of functional diversification through evolutionary processes. Nevertheless, conserved essential domains were preserved in all five APN paralogs (APN1-5), including the glutamine-activated zinc-binding motif (GAMEN) and the canonical zinc coordination triad (HEXXHX18E) (Fig. 2). This structural conservation suggests maintenance of core enzymatic functions despite sequence heterogeneity.

**Fig. 2.**
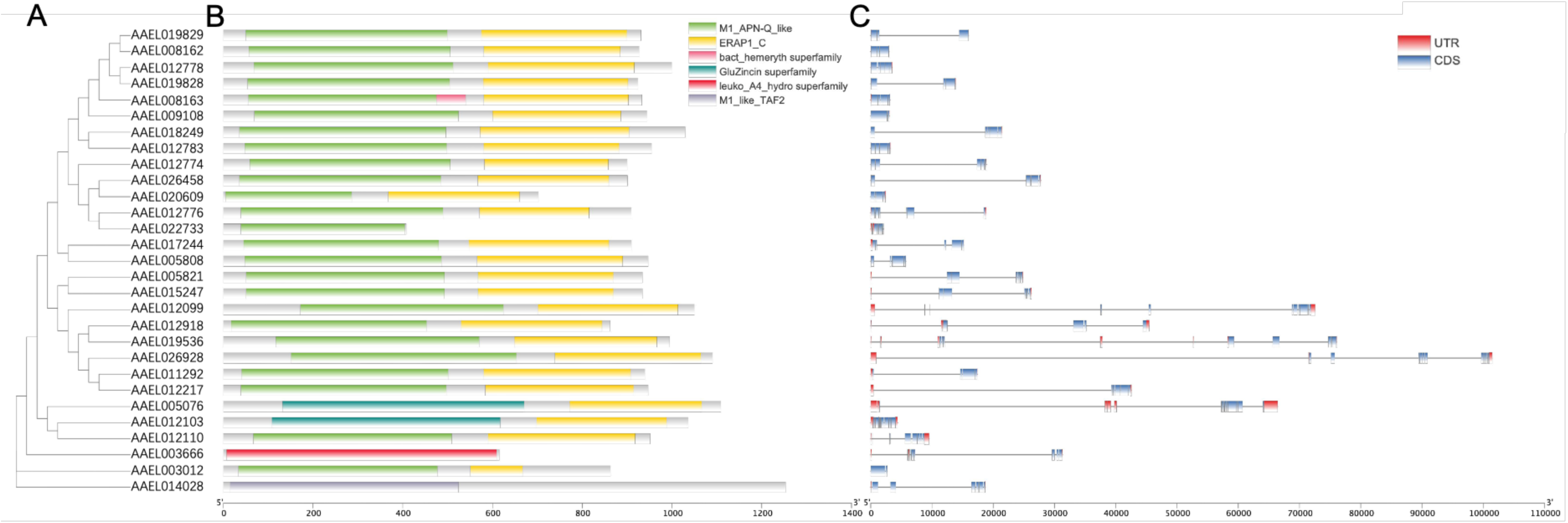
**The gene structure and conserved motif analyses of *AeAPN* genes**. (A) The phylogenetic relationships of AeAPN proteins. (B) Conserved motif distribution of AeAPN proteins. The different conserved motifs are represented by different colored boxes. (C) Gene structures of *AeAPN* genes. The UTR, CDS, and introns are indicated by red boxes, blue boxes, and black lines, respectively.

3.3 **Identification and Phylogenetic Analysis of AeAPNs**

To elucidate evolutionary dynamics within the AeAPN gene family, we performed a comparative phylogenetic analysis using full-length protein sequences from three medically important mosquito genera: *Ae. aegypti*, *Culex quinquefasciatus*, and *An. gambiae*. The circular phylogenetic tree revealed eight well-defined clades (Fig. 3), demonstrating conserved phylogenetic distribution patterns across these dipteran species. Notably, interspecific sequence homology consistently exceeded intraspecific homology levels, suggesting that the diversification events within the APN gene family predate the speciation processes of contemporary mosquito genera. Particularly significant was the phylogenetic segregation of Bti toxin-binding receptors (APN1-5), which occupied distinct clades with substantial branch length divergences (Fig. 3). This topological separation implies both significant divergence in secondary structural configurations and potential functional differentiation among these receptor subtypes.

**Fig. 3.**
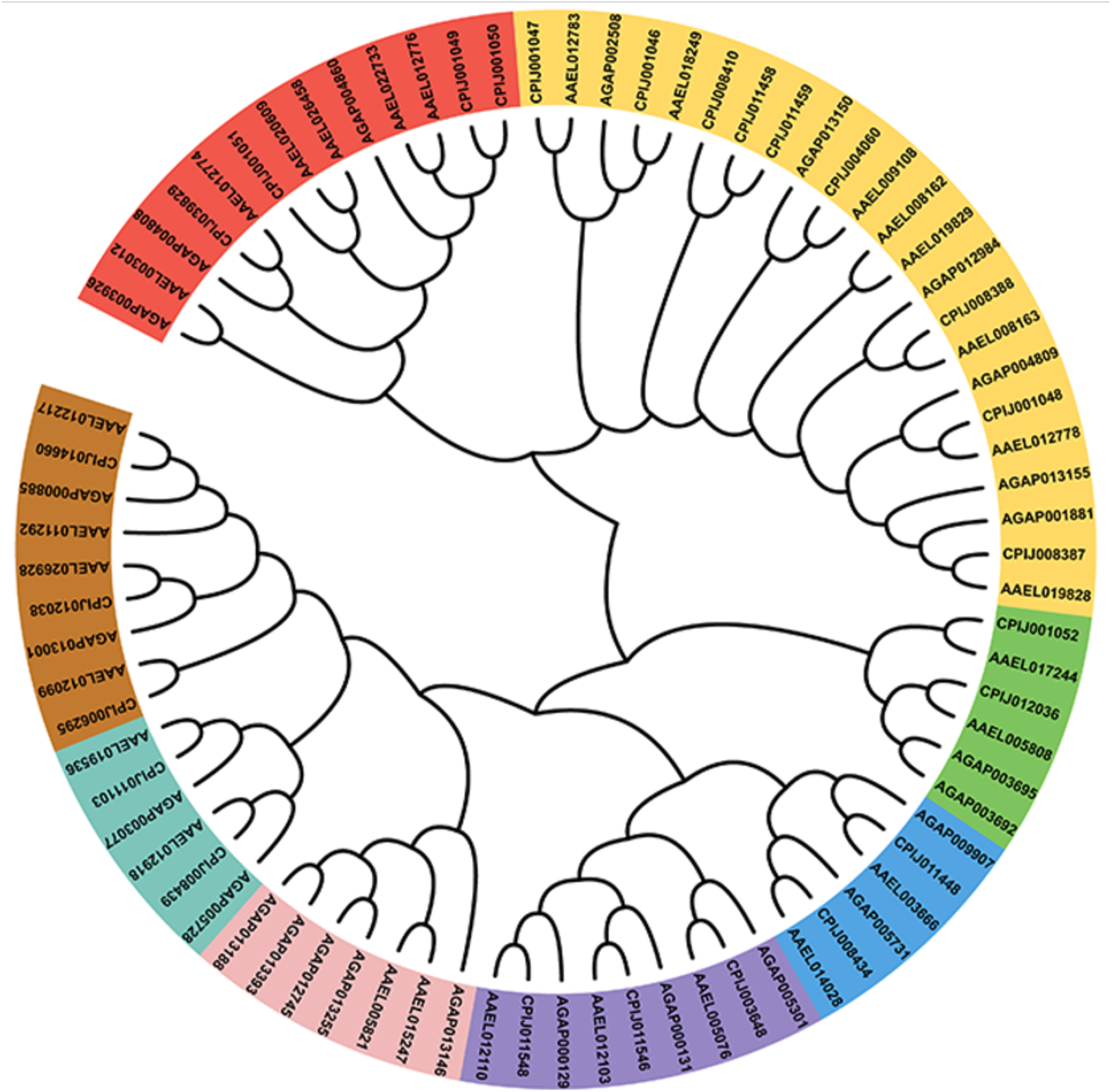
Phylogenetic analysis of APNs subunits from three mosquitoes. Phylogenetic tree constructed by using amino acid sequences of *Ae. aegypti*, *Cu. quinquefasciatus*, and *An. gambiae*. The APN proteins are clustered into 8 subgroups, marked by different colors.

### 3.4 Expression pattern analysis of AeAPN gene family

Transcriptomic quantification of *AeAPN* gene expression across developmental stages [1st- instar larvae (L1), 2nd-instar larvae (L2), 3rd-instar larvae (L3), 4th-instar larvae (L4), pupal stage, male adult, and female adult] and tissue compartments (intestinal and extra-intestinal tissues) revealed distinct spatial-temporal regulation patterns. Specifically, fifteen *AeAPN* genes demonstrated gut-specific expression, while nine *AeAPNs* exhibited peak transcriptional activity in the L3 stage, including APN1-3 and 5 (Fig. 4). These findings highlight the crucial role of APN as a digestive enzyme within the gut of larvae. These intestinal-enriched APNs, particularly APN1-3 and 5 that were previously identified as Bti-binding proteins, may serve as primary receptors mediating Bti toxin susceptibility in *Ae. aegypti*. Furthermore, three APN isoforms showed significant upregulation during pupal metamorphosis (Fig. 4), so they are potentially associated with tissue remodeling. Intriguingly, six male-biased and two female-biased APNs displayed adult stage-specific expression patterns (Fig. 4), suggesting their potential involvement in sex-specific physiological processes including post-eclosion development, nutrient metabolism, and xenobiotic detoxification.

**Fig. 4.**
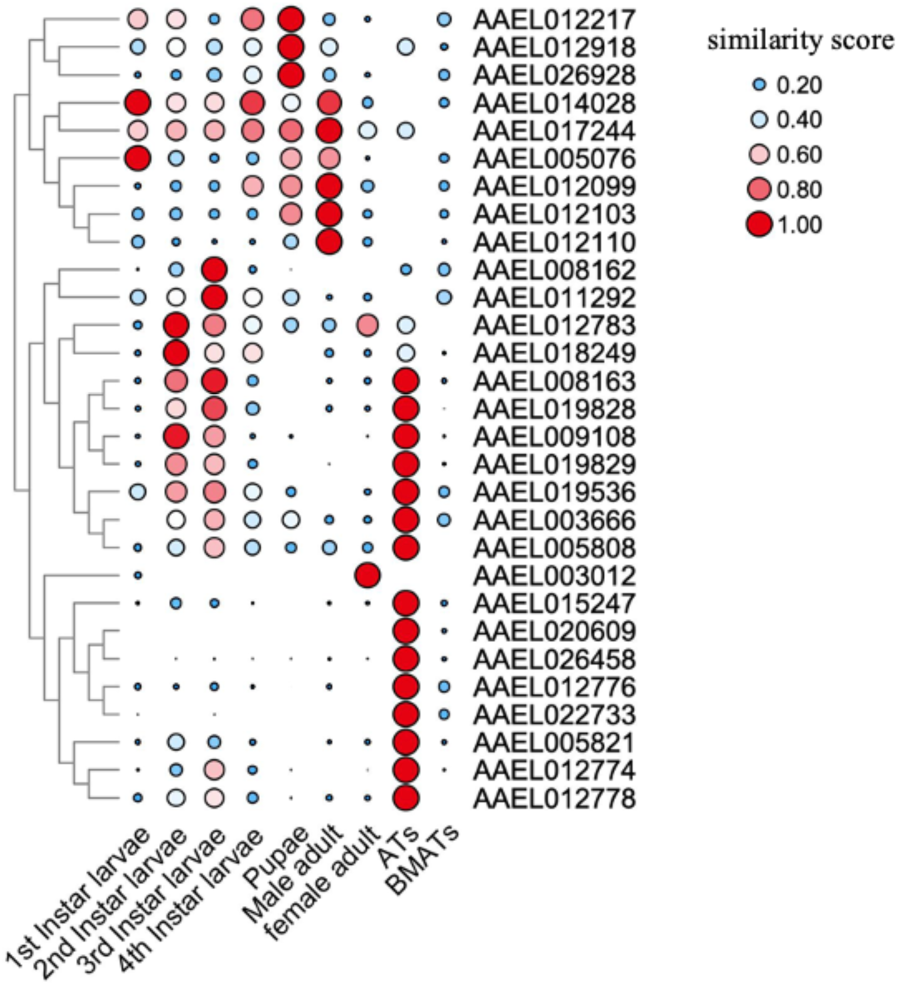
Spatial and temporal expression of APN genes in *Ae. aegypti*. Expression patterns of the *AeAPN* genes in the 1st-instar larvae (L1) to the 4th-instar larvae (L4), pupal stage, male adult, female adult, intestinal tissues (ATs), and extra-intestinal tissues (BMATs). Expression levels are shown based on bubble size and color.

### 3.5 Ligand blotting detection and ELISA binding assay of AeAPN3 protein

In our previous identification of Bti toxin binding proteins within the midgut epithelium of *Ae. aegypti* mosquitoes, we found that APN3 exhibits specific binding affinity to the Cry11Aa toxin. To systematically characterize the potential interaction between AeAPN3 and Cry4Ba toxin, we cloned the complete coding sequence (CDS) of AeAPN3 into a pColdTMGST prokaryotic expression vector (Fig. S1A) for expressing AeAPN3 protein in *E. coli* BL21(DE3) competent cells. Following nickel-affinity chromatography purification, the recombinant GST-tagged AeAPN3 protein was subjected to SDS-PAGE analysis, which confirmed the production of a predominant protein band with an apparent molecular mass of ∼128 kDa (Fig. S1B), consistent with the predicted molecular weight of the fusion protein. Western blot analysis using anti-GST monoclonal antibodies further validated the identity of the purified recombinant protein (Fig. S1C).

The interaction between AeAPN3 and Cry4Ba toxin was qualitatively analyzed through complementary ligand-binding methodologies. Ligand blot analysis validated specific binding bands corresponding to the APN3-GST fusion protein and Cry4Ba toxin, providing preliminary evidence for direct protein-protein interaction between the purified recombinant AeAPN3 and the Cry4Ba toxin (Fig. 5A). Furthermore, quantitative validation through the ELISA binding assay demonstrated a dissociation constant (Kd) of 20.53 nM, which suggested a high affinity between the Cry4Ba toxin and recombinant AeAPN3 protein (Fig. 5B). These results suggested that AeAPN3 is a putative receptor of Cry4Ba.

**Fig. 5.**
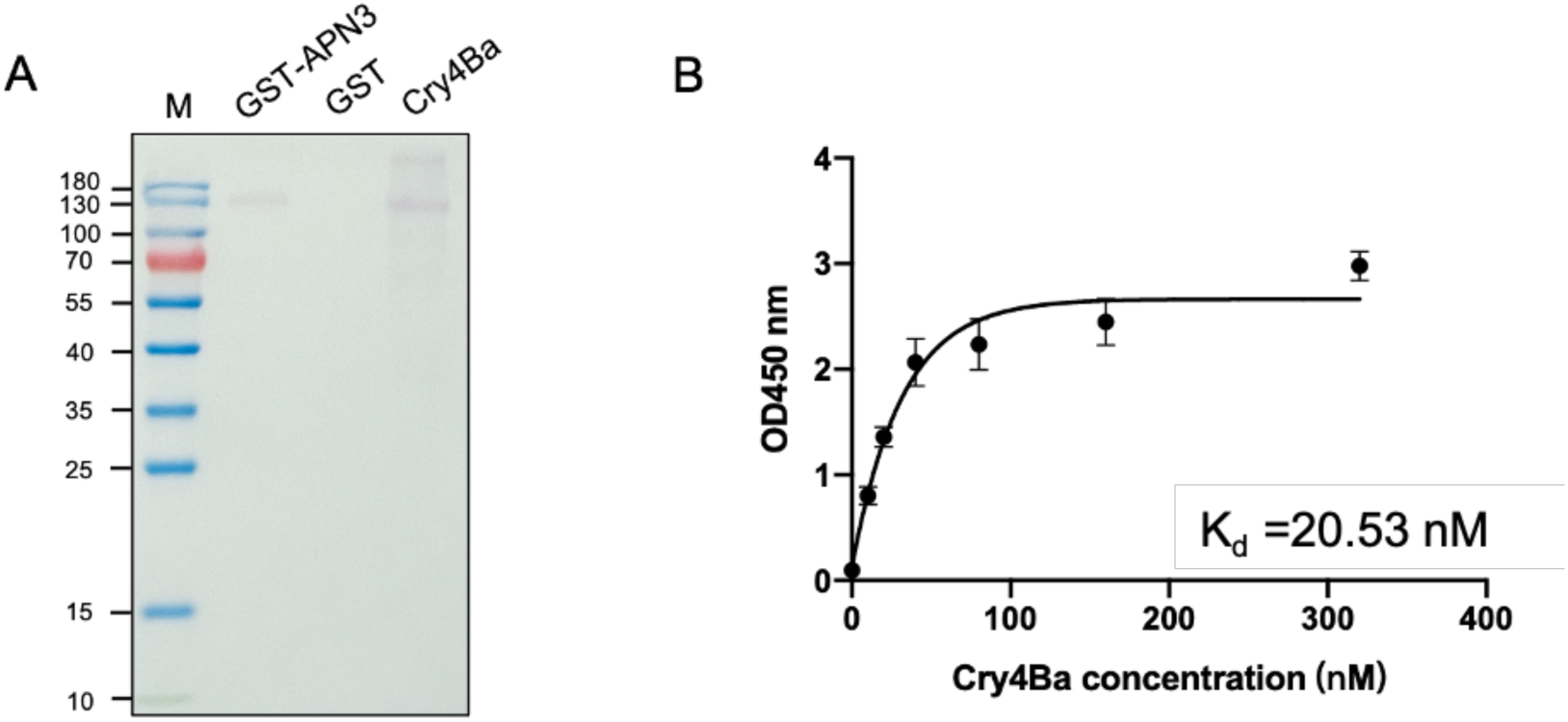
***In vitro* binding affinity of GST-APN3 protein to Cry4Ba toxin.** (A) Ligand blot analysis for the interaction between APN3 and Cry4Ba. (B) ELISA analysis for the binding affinity of APN3 to Cry4Ba.

### 3.6 Establishment of homozygous AeAPN3 knockout strain

To knockout AeAPN3, we injected a mixture of two sgRNAs targeting exon 3 of AeAPN3 and Cas9 protein into freshly laid eggs from an EXU-Cas9 knock-in strain. A total of 200 fresh eggs were consecutively injected, with only approximately 6% (12/200) successfully hatching, and among them, 66.7% (8/12) larvae eventually developed to adults (Table S3). However, successful site-specific mutations within AeAPN3 were identified in all surviving G0 mosquitoes (8/8) by sequencing PCR products from individual DNA samples (Table S3).

The homozygous knockout (KO) *Ae. aegypti* strain for *Ae*APN3 (named *Ae*APN3-KO-EXU) was generated using the reverse genetics approach. *Ae*APN3-KO-EXU presented a 100 bp deletion and 2 bp insertion between two of the CRISPR/Cas9 target sites (Fig. 6), resulting in truncated protein production (Fig. S2). To eliminate the influence from endogenous expression of the CRISPR/Cas9 system and *kmo* gene deletion on subsequent functional verification, red fluorescent and white eye individuals were removed from *Ae*APN3-KO-EXU strains by fluorescent screening and gene sequencing. Consequently, a new homozygous knockout strain for AeAPN3 named as AeAPN3-KO was established by selecting individuals with non-fluorescent normal eyes.

**Fig. 6.**
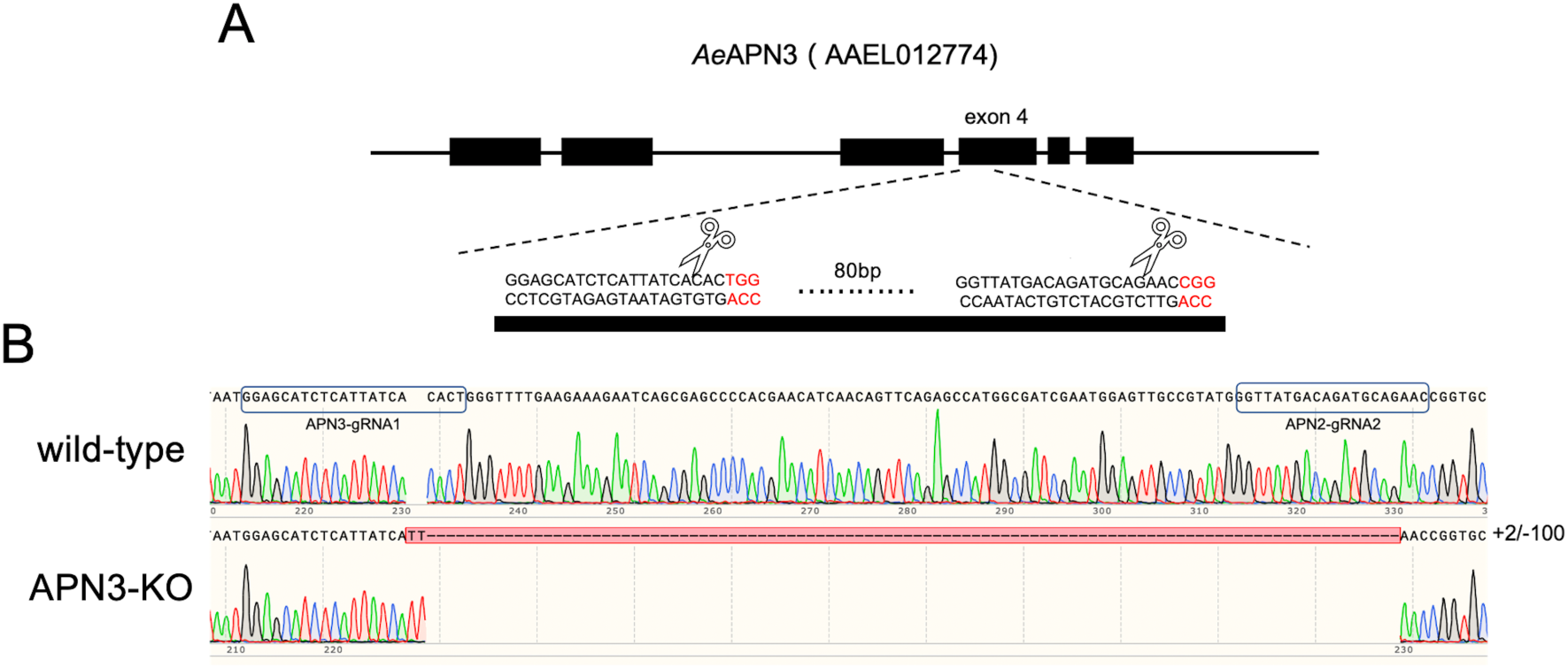
Generation of APN1 knockout *Ae. Aegypti* strains. (A) Schematic representations of the APN1 locus, with the third exon contain 20-nucleotide sgRNA target sequences, and the PAM sequence shown in red. (B) Aligned Sanger-sequencing trace of PCR-amplified genomic DNA from wild-type and APN1-KO strains with specific primers (Table S1) spanning the genomic RNA targeted region.

### 3.7 Resistance to Cry4Ba protoxin caused by AeAPN3 knockout

Susceptibility to the Cry4Ba and Cry11Aa protoxins using six gradient concentrations were tested in the *Ae*APN3-KO strain with the original susceptible wild-type strain as a negative control. Bioassays results showed a LC50 value of 7.275 (95%CI: 6.672-7.986) μg/ml to the Cry4Ba protoxin for *Ae*APN3-KO, which was approximately 2.938- to 4.108-fold higher than the susceptible wild-type strain (1.771 (95%CI: 1.663–1.888) μg/ml] (Table 1). This result indicated that AeAPN3 is a functional receptor for Cry4Ba. However, for bioassays of Cry11Aa protoxin, the LC50 value of the *Ae*APN3-KO strain (0.521 (95%CI: 0.473-0.568) μg/ml] was lower than wild-type strain (0.602 (95%CI: 0.526–0.685) μg/ml], but not significantly different (Table 1).

**Table 1.** Susceptibility of *Ae. aegypti* strains to Cry4Ba and Cry11Aa toxins.

| Toxin | Strains | n | Slope (SE) | LC50 ($\mu\text{g/mL}$) (95% CI) | RR |
| --- | --- | --- | --- | --- | --- |
| Cry4Ba | wild-type-1 | 1125 | 2.947 (0.098) | 1.771 (1.663–1.888) | 1 |
|  | wild-type-2 | 1350 | 3.023(0.159) | 2.476 (1.753-3.175) | 1.398 |
|  | APN3-KO | 1350 | 2.953 (0.142) | 7.275 (6.672-7.986) | 2.938-4.108 |
| Cry11Aa | wild-type-1 | 1125 | 1.747 (0.110) | 0.602 (0.526–0.685) | 1 |
|  | wild-type-2 | 1350 | 2.251 (0.12) | 0.801 (0.729-0.876) | 1.331 |
|  | APN3-KO | 1350 | 2.291 (0.122) | 0.521 (0.473-0.568) | 0.650-0.865 |

To elucidate the role of AeAPN3 in response to Cry4Ba, we conducted a comparative analysis of mortality rates between the AeAPN3-KO and wild-type strains. Our results demonstrate that the mortality rate of AeAPN3-KO was significantly lower than wild-type across all tested Cry4Ba concentrations (*P* < 0.0001, t-test). Specifically, at LC50 concentration (2.5 μg/mL) for wild-type, the mortality rate for AeAPN3-KO was only 7.1±5.3%. Furthermore, at a diagnostic dose (10 μg/mL) of Cry4Ba for wild-type, the mortality rate for AeAPN3-KO increased to 58.7±12.2% (Fig. 7). Notably, when Cry4Ba concentration was further elevated to 18.75 μg/mL, there was a corresponding increase in mortality rate observed in APN3-KO mosquitoes, reaching 88.4±4.4%. These findings indicated that APN3 plays a crucial role in mediating the activity of Cry4Ba at low lethal concentrations (Fig. 7).

**Fig. 7.**
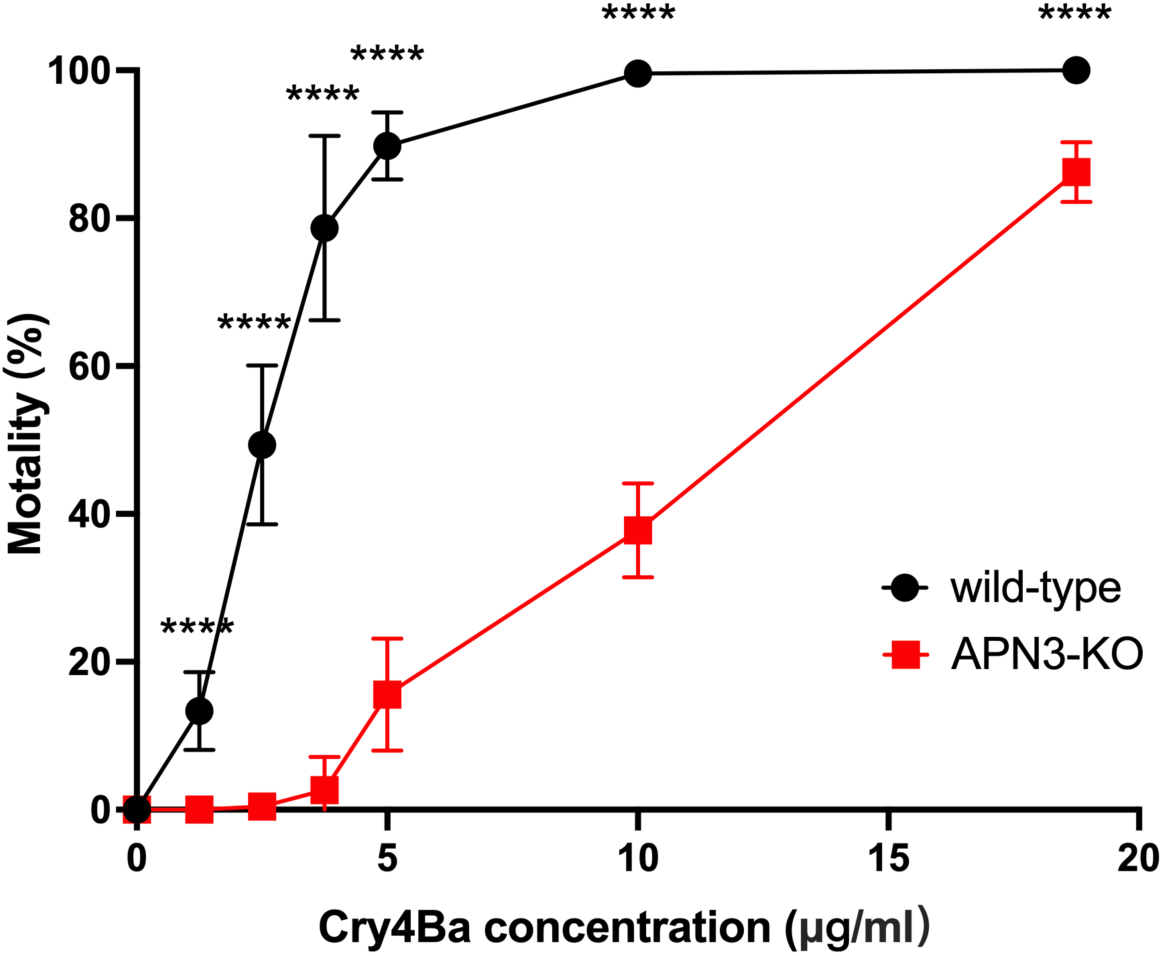
R**e**sponse **of the APN3-KO and wild-type strains to Cry4Ba toxin.** Twenty-five larvae from wild-type and APN3-KO strains were placed in cups containing six different concentrations of Cry4Ba toxin. Larval mortality was assessed after 24 hours. Each experimental setup included three replicates, and three independent experiments were performed in total. Each point represents the mean ± SD from three biological replicates. Differences between groups were analyzed by t-test (****, *P* < 0.0001).

## 4 Discussion

As a class of proteolytic enzymes, APN (EC3.4.11) is primarily distributed in the intestinal and salivary glands of insects, playing a crucial role in protein and peptide digestion by effectively cleaving N-terminal amino acid residues^29^. Eight to ten APN genes have been found in the genomes of Lepidopteran insects such as *Bombyx mori*, *Helicoverpa armigera*, *Plutella xylostella*, and *Spodoptera frugiperda*, respectively^29^. Additionally, five APN transcripts were identified in the transcriptome in the *Nilaparvata lugens* of Hemiptera^30^, while eleven APN transcripts were discovered in the midgut transcriptome from *Chrysomela populi* of Coleoptera^31^. Notably, twenty-nine, twenty-five, and twenty-five APN genes were respectively exhibited in the genomes of three major pathogenic mosquitoes (*Ae. aegypti*, *Cu. quinquefasciatus*, and *An. gambiae*). The present finding suggests a pronounced expansion trend of the APN gene family in Diptera mosquitoes compared to other insect orders (Fig. 2 and 3). However, analysis of the spatiotemporal transcriptome of *Ae. aegypti* revealed that at least 10 APN transcripts exhibited negligible or very low expression levels throughout their life cycle, suggesting the presence of redundant copies and nonfunctional pseudogenes within the *Ae. aegypti* APN family during evolution (Fig. 2 and 3).

By analyzing the spatiotemporal transcriptome of *Ae. aegypti*, our investigation identified that 10 APN transcripts remained constitutively silent or exhibited minimal expression throughout ontogenetic development, suggesting that the APN family in *Ae. aegypti* may have undergone duplication events resulting in nonfunctional pseudogenes (Fig. 4). Through physical mapping, we observed that while *Ae. aegypti* APNs were distributed across three chromosomes, they predominantly clustered in the upper and lower regions of chromosome 1 (Fig. 4). This clustering pattern was also evident in Lepidopteran APNs, such as APN1 and APN3, APN2, APN4 and APN6, and APN7 and APN9 are arranged adjacent to each other on the genome of *He. armigera*, indicating that gene duplication events of APNs mainly occur in close proximity^32^. Through the investigation of genetic evolution in five Lepidoptera species, it was discovered that there exists collinearity among APN genes across these different orders. The arrangement and orientation of APN gene groups or species within the genome exhibit a general consistency, indicating a conservation of the APN gene family during evolutionary processes in Lepidoptera^32^. However, the genetic evolutionary patterns of mosquito APNs show significant differences in *Ae. aegypti*. While *AeAPN1*, *AeAPN3*, and AAEL012776 are closely located on the chromosome, they exhibited substantial variations in gene structure and belong to two different major branches, reflecting the diversity in expansion evolution of the mosquito APN gene family (Fig. 1-3).

The APN protein is predominantly expressed as both a membrane-bound and soluble form. In the insect gut, the membrane-bound APN protein attaches to the brush border membrane of epithelial cells through its GPI anchor site at the C terminus. In *Ae. aegypti*, 11 APNs were predicted to have GPI anchor sites, including Bti-binding proteins APN1, APN2, APN4, and

APN5 (Tables S2), using three different GPI prediction software tools (big-PI Predictor, PredGPI predictor, and KohGPI). However, Chen *et al*. identified a potential GPI anchor site on APN3 using DGPI software, which suggests variations in prediction results among different algorithms. Thus, further verification is required through cell sublocalization experiments^15^. In mammals, GPI-APNs had been identified as membrane receptors for various pathogens^33–36^. Conversely, in insects, GPI-APNs were primarily acknowledged as crucial receptors for Bt Cry toxins and played significant roles in both toxicological responses and resistance mechanisms against Bt toxins^18, 22^. Furthermore, it was discovered that APN functions as a receptor protein for pea enation mosaic virus (PEMV) and lectin proteins in pea aphids (*Acyrthosiphon pisum*)^37,38^.

This study revealed the significance of glycosylation sites in APN for its binding process with Cry proteins. In Bti-binding proteins APN1-5 of *Ae. aegypti*, a number of O-glycosylation and N-glycosylation sites were predicted (Table S2). However, it should be noted that glycosylation is not necessarily the key mediator of the interaction between APN and Cry toxins. Several insect APNs expressed in prokaryotic systems also exhibited specific binding abilities to Cry toxins. For instance, fragments Ile135-Pro198 and Ile135-Gly174 from BmAPN1 expressed in a prokaryotic system retained full binding ability to Cry1Aa, while fragment Gly155-Pro198 retained partial binding capacity^39^. Although AeAPN3 was found to specifically bind to Cry11Aa through GST-Pull down and Co-IP experiments, it was not identified as a Cry4Ba- binding protein through mass spectrometry analysis^14^. To further investigate the interaction between AeAPN3 and Cry4Ba, we cloned the full-length *AeAPN3* gene into the PET32a vector for prokaryotic expression. Following IPTG induction, His-tagged AeAPN3 protein predominantly formed inclusion bodies within precipitates. To enhance solubility of AeAPN3 protein, we introduced AeAPN3 into the pCold-GST vector for expression. Results revealed that GST-tagged APN3 was present both in supernatant and precipitate fractions. The soluble form of AeAPN3 with a GST tag was purified using a GST chromatography column (Fig. S1). Ligand blotting and ELISA further confirmed a strong interaction between APN3 and Cry4Ba with an affinity of 20.53 nM. This indicated that APN3 is a membrane-bound protein for both Cry4Ba and Cry11Aa, while suggesting that their interaction may not rely on glycosylation modifications (Fig. 5). Some studies observed similar results in interactions between AeAPNs such as AeAPN1^15^ or AeAPN2a^16^ with Bt toxins.

Although APNs have been confirmed as the major binding proteins for Bt toxins in numerous insect species, it should be noted that not all APNs are capable of binding to Cry toxins. Even if APN proteins could bind with Cry toxins, it does not necessarily mean that they are functional receptors mediating Cry activity. In *He. armigera*, ligand blot analysis revealed that HaAPN1 could bind to Cr1A, and overexpression of HaAPN1 in Sf21 insect cells caused aberrant cell morphology^17^. However, CRISPR/Cas9-mediated knockout of *HaAPN1*, *HaAPN2*, and *HaAPN5* did not affect the sensitivity of larvae to Cry1A and Cry2A, likely excluding a major contribution of these genes to Bt toxicity mechanisms^23^. Similarly, GST-Pull down and Co-IP experiments showed that *AeAPN1* could specifically bind to Cry4Ba and Cry11Aa, while *AeAPN2* could specifically bind to Cry11Aa in *Ae. aegypti*^14^. Nevertheless, subsequent bioassays conducted on both wild-type strains and CRISPR/Cas9-generated knockout strains lacking either *AeAPN1* or *AeAPN2* showed no change in sensitivity towards these two types of Cry toxins^22^. These findings suggest that neither APN1 nor APN2 serve as essential functional receptors for Bti^22^. However, knocking out *AeAPN3* resulted in approximately 4-fold relative resistance to Cry4Ba despite having no significant effect on sensitivity towards Cry11Aa. This indicated the potential involvement of AeAPN3 in mediating Cry4Ba functionality.

The mechanism of Bt action is highly intricate and may involve the mediation of multiple membrane-bound receptors. In the diamondback moth, knockout of either *PxABCC2* or *PxABCC3* did not result in significant Cry1Ac resistance, whereas simultaneous knockout of both genes led to a remarkable 8000-fold increase in Cry1Ac resistance in larvae^40, 41^. These findings indicate the redundant role of *PxABCC2* and *PxABCC3* as functional receptor proteins mediating Cry1Ac activity^40,41^. In *Ae. aegypti*, simultaneous knockout of *AeAPN1* and *AeAPN2* had no impact on sensitivity to Cry4Ba and Cry11Aa^22^. However, results from AeAPN3-KO experiments suggest that *AeAPN3* played a more prominent role in the toxic mechanism of Cry at low concentration. At high concentration, Cry toxins may exert its toxicity through multiple receptors or pathways (Fig. 7). In the classical Bt model, activated Cry toxins first forms oligomers by interacting with CAD before binding GPI-anchored proteins (APN and ALP) on the cell membrane surface causing perforation leading to cell death^42^. The role of APN may be similar to that of a “magnet”, absorbing free Cry onto the cell membrane, subsequently enriched Cry binds with other functional receptors (such as ABC transporters) to complete the perforation process. Therefore, low doses of Cry require mediation by binding receptors like APN. However, once the concentration reaches a certain threshold level, direct interaction between Cry and other functional receptors can occur without APN absorption.

In addition, there was a significant association between APN and Bt resistance mechanisms. For instance, in the Cry1Ac high-resistance strain of *He. armigera* (BT-R), it was discovered that the O-glycosylation site-enriched region of APN1 lacked 22 amino acids, resulting in the loss of binding affinity with Cry1Ac^43^. Similarly, in the highly resistant Cry1Ac strain of fall armyworm (GLEN-Cry1Ac-BCS), a significant down-regulation in APN1 expression level was observed while APN6 expression level showed a notable up-regulation^44^. Further genetic linkage analysis revealed that Cry1Ac resistance was linked to the down-regulation of APN1, while the up-regulation of APN6 expression may compensate for the loss-of-function caused by this down-regulation, thereby minimizing fitness costs^44^. Consequently, further research may investigate whether other *Ae. aegypti* APNs affect Bt activity through alternative mechanisms and if alterations in enzymatic activity directly contribute to Bt resistance.

## 5 Conclusions

This study demonstrated through ligand blot and ELISA analyses that *Ae*APN3 functions as a high-affinity binding receptor for the Cry4Ba toxin. Gene knockout experiments further confirmed that *AeAPN3* depletion significantly enhanced resistance to Cry4Ba toxin, thereby establishing *Ae*APN3 as the functional receptor mediating Cry4Ba toxicity.

## Acknowledgements

This project supported by the Science Foundation of Fujian Province, China grant number 2023J011576 to JW.

## Supplemental Information

**Fig. S1.**
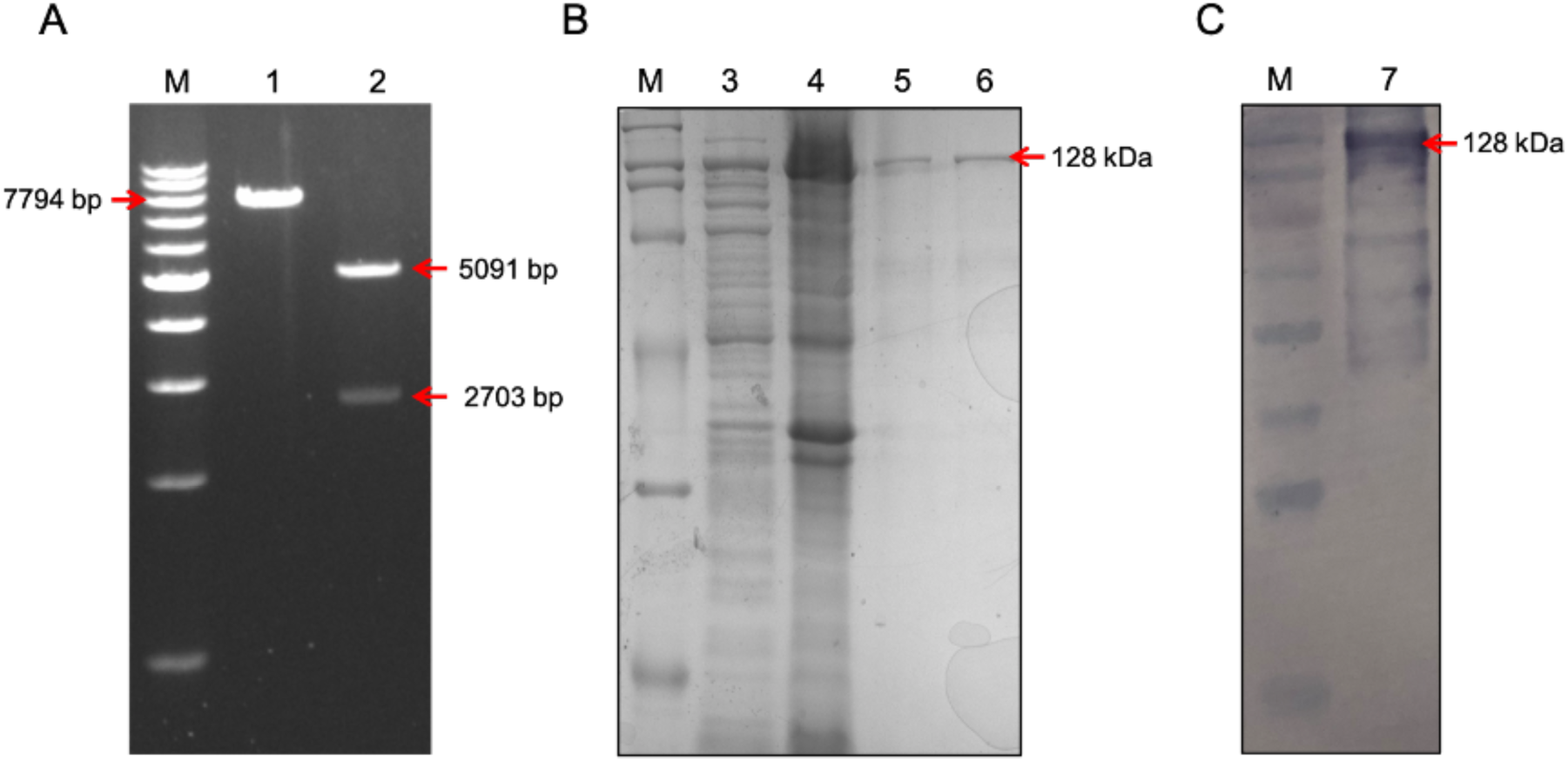
Prokaryotic expression of GST-APN3 protein. (A) Single and double digestion of GST- APN3 vector. Lane M, 1 kb DNA Marker, Lane 1: GST-APN3 vector digested by XhoI, Lane 2: GST- APN3 vector digested by XhoI and HindIII. (B) Prokaryotic expression and purification of GST-APN3 protein. Lane M: Protein Molecular Weight Marker (Broad), Lane 3: supernatant of degree induction with IPTG, Lane 4, precipitate of degree induction with IPTG, Lane 5 and 6: purified GST-APN3 proteins. (C) Western blot analysis of the expression of GST-APN3 protein. Lane M: Prestained Color Protein Marker, Lane 7: identification of GST-APN3 protein using GST-Tag monoclonal antibody.

**Fig. S2.**
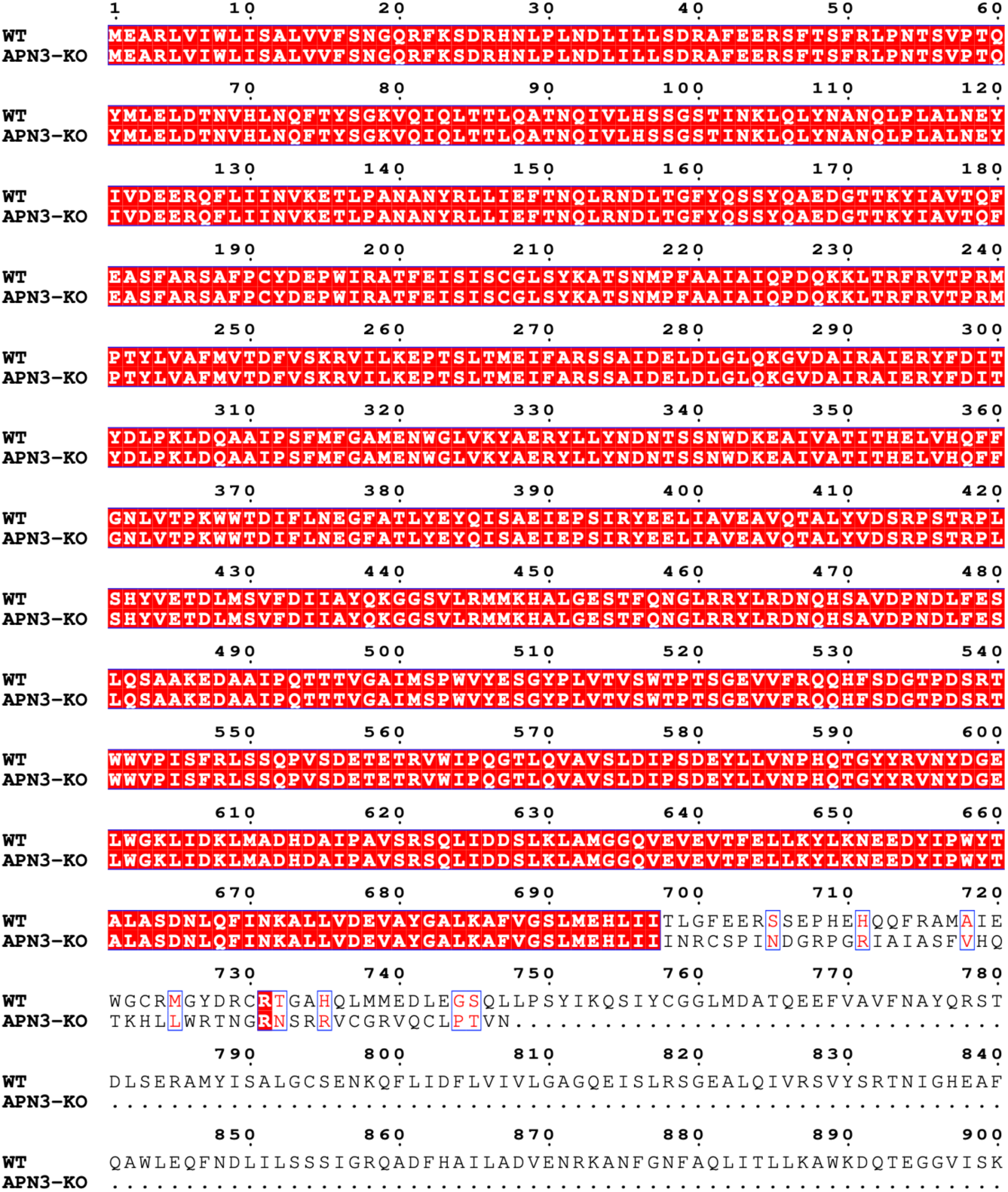
Amino acid sequence alignment of APN3 isoform from wild-type and APN3-KO strains. Red highlights and ed text (with blue box) mean the same and similar amino acid sequence between wild-type and APN3-KO strains, respectively. The dots indicate premature termination of translation, which prevents the formation of complete amino acid sequences after knockout the APN3 gene.

**Table S1.**
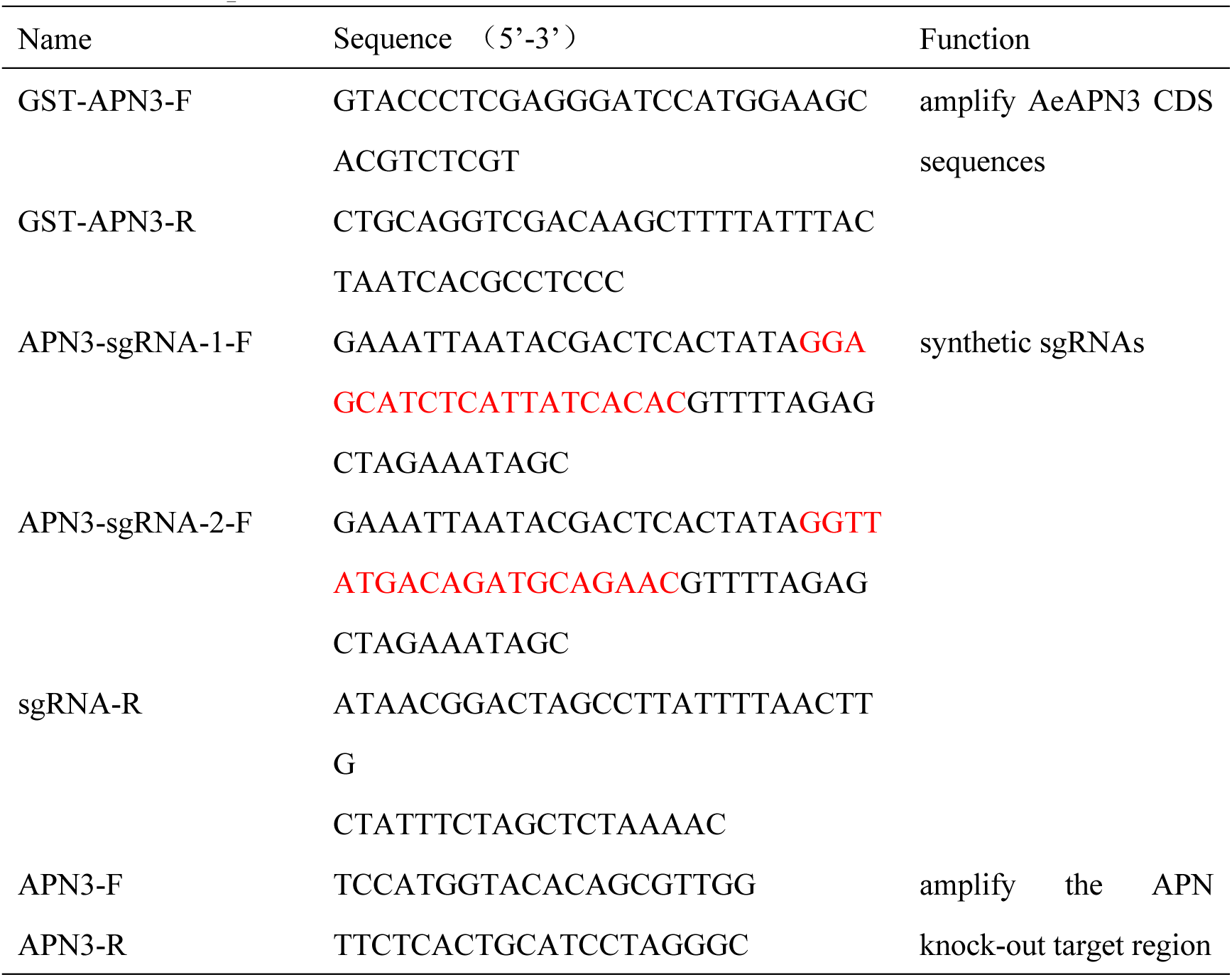
PCR primers.

**Table S2.** Gene feature profiling of APNs in *Ae. Aegypti*.

| Gene ID | Gene name | big-PI Predictor | PredGPI | KohGPI | Signal IP | N-glycosylated | O-glycosylated |
| --- | --- | --- | --- | --- | --- | --- | --- |
| AAEL003012 | glutamyl aminopeptidase | N/A | N/A | N/A | ● | 9 | N/A |
| AAEL003666 | leukotriene A-4 hydrolase | N/A | N/A | N/A | N/A | 2 | 2 |
| AAEL005076 | endoplasmic reticulum aminopeptidase 2 | N/A | N/A | N/A | N/A | 15 | 24 |
| AAEL005808 | aminopeptidase N | ● | ● | ● | ● | 2 | 9 |
| AAEL005821 | membrane alanyl aminopeptidase | ● | ● | ● | ● | 3 | 4 |
| AAEL008162 | aminopeptidase N | N/A | N/A | N/A | ● | N/A | 6 |
| AAEL008163 | aminopeptidase N | N/A | N/A | N/A | N/A | 1 | 6 |
| AAEL009108 | aminopeptidase N | N/A | ● | ● | ● | 3 | 7 |
| AAEL011292 | aminopeptidase N | N/A | N/A | N/A | ● | 8 | 1 |
| AAEL012099 | aminopeptidase N | N/A | N/A | N/A | N/A | 6 | 12 |
| AAEL012103 | aminopeptidase N | N/A | N/A | N/A | N/A | 13 | N/A |
| AAEL012110 | aminopeptidase N | N/A | N/A | N/A | N/A | 9 | 3 |
| AAEL012217 | aminopeptidase N | N/A | N/A | N/A | ● | 5 | 4 |
| AAEL012774 | aminopeptidase N | N/A | N/A | N/A | ● | 2 | 6 |
| AAEL012776 | aminopeptidase N | N/A | ● | N/A | ● | 3 | 5 |
| AAEL012778 | aminopeptidase N | ● | ● | ● | ● | 1 | 0 |
| AAEL012783 | aminopeptidase N | ● | ● | ● | ● | 8 | 6 |
| AAEL012918 | puromycin-sensitive aminopeptidase | N/A | N/A | N/A | N/A | 3 | 26 |
| AAEL014028 | transcription initiation factor TFIID subunit 2 | N/A | N/A | N/A | N/A | N/A | 42 |
| AAEL015247 | membrane alanyl aminopeptidase | ● | ● | ● | ● | 4 | 4 |
| AAEL017244 | aminopeptidase N | N/A | ● | N/A | ● | 9 | 4 |
| AAEL018249 | aminopeptidase N | ● | ● | ● | N/A | 11 | 44 |
| AAEL019536 | glutamyl aminopeptidase | N/A | N/A | N/A | N/A | 7 | 13 |
| AAEL019828 | aminopeptidase N | N/A | ● | ● | ● | 2 | N/A |
| AAEL019829 | glutamyl aminopeptidase-like | N/A | N/A | ● | ● | 5 | 2 |
| AAEL020609 | aminopeptidase N | N/A | N/A | N/A | N/A | 3 | 1 |
| AAEL022733 | endoplasmic reticulum aminopeptidase 1 | N/A | N/A | N/A | ● | 1 | 1 |
| AAEL026458 | aminopeptidase N | N/A | N/A | N/A | ● | 6 | 1 |
| AAEL026928 | aminopeptidase N | N/A | N/A | N/A | N/A | 12 | 26 |

**Table S3.** Injection, survival, and mutagenesis rates mediated by CRISPR/Cas9 constructs in the ExuCas9-*kmo* strain.

| Injected Strain | Injected Component | Injected G0 Embryos | G0 Hatch/Survivors | G0 Mutants |
| --- | --- | --- | --- | --- |
| ExuCas9-*kmo* | Cas9 protein, APN3-sgRNA1,<br>APN3-sgRNA2 | 200 | 12/8 | 8 |

